# MicroCT-Based Imaging of Microvasculature in Bone and Peri-Implant Tissues

**DOI:** 10.1101/2023.03.08.531678

**Authors:** David Haberthür, Oleksiy-Zakhar Khoma, Tim Hoessly, Eugenio Zoni, Marianna Kruithof-de Julio, Stewart D. Ryan, Myriam Grunewald, Benjamin Bellón, Rebecca Sandgren, Stephan Handschuh, Benjamin E. Pippenger, Dieter Bosshardt, Valentin Djonov, Ruslan Hlushchuk

## Abstract

Angiogenesis is essential for skeletal development, bone healing, and regeneration. Improved non-destructive, three-dimensional (3D) imaging of the vasculature within bone tissue benefits many research areas, especially implantology and tissue engineering.

X-ray microcomputed tomography (microCT) is a well-suited non-destructive 3D imaging technique for bone morphology. For microCT-based detection of vessels, it is paramount to use contrast enhancement. Limited differences in radiopacity between perfusion agents and mineralized bone make their distinct segmentation problematic and have been a major drawback of this approach. A decalcification step resolves this issue but inhibits the simultaneous assessment of bone microstructure and vascular morphology. The problem of contrasting becomes further complicated in samples with metal implants.

This study describes contrast-enhanced microCT-based visualization of vasculature within bone tissue in small and large animal models, also in the vicinity of the metal implants. We present simultaneous microvascular and bone imaging in murine tibia, a murine bone metastatic model, the pulp chamber, gingiva, and periodontal ligaments. In a large animal model (minipig), we perform visualization and segmentation of different tissue types and vessels in the hemimandible containing metal implants. We further demonstrate the potential of dual-energy imaging in distinguishing bone tissue from the applied contrast agents.

This work introduces a non-destructive approach for 3D imaging of vasculature within soft and hard tissues near metal implants in a large animal model.

## Introduction

Angiogenesis, the formation of new blood vessels from preexisting vessels, is crucial for skeletal development and bone healing and regeneration [5,10,42,44,52]. In addition to carrying nutrients and growth factors, these newly formed blood vessels serve as a delivery route for stem cells and progenitor cells to the bone defect site [13,29,30]. In the case of bone grafts, many synthetic bone grafts fail to bridge critically sized defects due to their inability to promote vascularization [33,42]. The structural nature of skeletal tissue makes three-dimensional (3D) imaging of its vasculature extremely difficult. Histology, a destructive and two-dimensional approach, remains the gold standard for assessing vasculature in bones [37].

Classic soft tissue imaging techniques such as light sheet microscopy or confocal laser scanning microscopy face challenges in their application due to the encasement of blood vessels in calcified tissue [37]. Despite recent advances in tissue-clearing-based imaging methods for craniofacial and other bones [3,34,43] and the whole body [4], the application of such methods remains challenging and is largely limited to small animal models. Simultaneously imaging both vasculature and bone tissue non-destructively in 3D has long been a challenge, especially in large bone grafts or near metal implants [35,42]. Thus, many research areas, such as tissue engineering, implantology, reconstructive surgery, bone biology, and bone metastatic disease, benefit from improved three-dimensional imaging of the vasculature within bone tissue.

In the last decades, X-ray microcomputed tomography (microCT) gained recognition as a non-destructive 3D imaging technique for bone morphology [17]. Due to the inherently low difference in X-ray absorption levels between vessels and different soft tissues, it is not easily feasible to distinguish such structures within the bone. To unambiguously detect vasculature within bone, it is necessary to instill the vessels with a contrast agent or use a casting method to fill the blood vessels. Another alternative to detect vessels within bone is synchrotron-radiation-based tomography, specifically employing phase-contrast tomography [12]. Although it has potential, synchrotron-radiation-based tomography is not as easily attainable for most preclinical researchers as microCT and has severe drawbacks in terms of visualized sample volume.

Currently, existing protocols for imaging the vasculature within bone via vascular replicas have drawbacks, such as disjoint vascular components or completely missing vascular segments [37,47]. The minor contrast difference between the perfusion agent used to generate the vascular network replica and the mineralized bone makes it problematic to distinguish bone tissue and vasculature. To enable proper visualization and segmentation of vasculature within bone tissue, decalcifying the bone samples has become practically a standard method [37,44]. This decalcification procedure makes simultaneous assessment of bone microstructure and vascular morphology impossible [37]. However, in a recent study by Rosenblum et al. [45], in a small animal model study without biomedical implants, this limitation was shown to be overcome by iterative microCT imaging (pre- and post-decalcification).

Intravascular contrast-agent-enhanced microCT has the potential to overcome the mentioned issues. It has become a method of choice for the evaluation of angiogenesis in bone tissue engineering and remodeling applications [42,44,55]. Barium sulfate and Microfil are the two most commonly used contrast agents in studies on the vascularization of bone tissue. Only some selected studies [44] managed to show better perfusion and thus visualization when perfusing the vasculature with barium sulfate. Previous studies have reported disadvantages associated with barium sulfate suspensions, including higher viscosity, which can result in incomplete vascular filling and weak or inhomogeneous signals, particularly in higher-resolution scans [44]. These issues may be attributed to particle aggregation, as indicated by various studies [27,31,40]. Although Microfil has probably been applied in a larger number of vasculature imaging studies than barium sulfate, Microfil was reported to have disadvantages like vascular damage [23] and poor or incomplete filling of the vasculature [31,40,44].

In the study of tumor models [19], assessing and non-destructively imaging tumor vasculature in 3D is a demanding task. Due to its intraosseous location, imaging is even more challenging in a bone metastatic disease model. Changes in bone and vasculature are of crucial importance for the progression of bone metastatic disease. Therefore, simultaneous imaging of bone and vasculature is highly desired, making a decalcification step unfavorable. It is therefore of paramount importance to develop an imaging method for assessing tumor vasculature in bone without decalcification, which renders the bone microstructure X-ray transparent. In this manuscript, we present such a method for visualizing intratumoral vasculature and defects in mineralized bone tissue.

The craniofacial skeleton, including the mandible, is a frequent location for distraction osteogenesis and other forms of bone regeneration and repair, for which vascularization is important [10,26]. Even though metal implants have revolutionized the treatment of patients with missing teeth or injured joints and bones [41,54], they are problematic for 3D imaging. The problem of imaging the vasculature is further compounded by their presence, for example, when studying vascularization in a jaw with metal implants. Due to their high X-ray absorption, metal implants produce beam hardening, partial volume, and low signal artifacts in the resulting tomographic datasets [24]. Interactions occurring at the tissue-implant interface are widely believed to play a crucial role in the success of implant placement and healing [1,54]. High-resolution tomographic imaging of the tissue-implant interface is complicated by artifacts from the metal implants in the resulting datasets. Imaging with increased acceleration voltage and at low resolutions alleviates these imaging artifacts. MicroCT imaging is the only available approach for non-destructively investigating the intact bone-implant interface in 3D [54] and in proximity to the implant surface (i.e. closer than 100 μm).

Implantology and osteology studies are often conducted in large animal models making the corresponding imaging even more complicated due to the lack of transgenic lines and the large size of the harvested samples. The Göttingen Minipig is widely recognized as a valuable large animal model in preclinical dental and orofacial research, mainly because of its anatomical similarities to humans [2,38,51]. Its bone structure and bone remodeling processes closely resemble those of humans, further enhancing its suitability for such studies, including ours.

The present study introduces a technique for microCT-based visualization of microvasculature within bone tissue in small and large animal models. Moreover, we demonstrate that this approach is suitable for simultaneous imaging and subsequent analysis of peri-implant hard and soft tissues, as well as their vascularization, in the vicinity of metal implants in a large animal model.

## Materials, Methods, and Results

### Animals

In this study, we used one 21-month-old transgenic VEGF male mouse (see [14] for more details), five 10-week-old CB17SCID male mice, three 12-week-old C57BL/6 mice, one 73-week-old C57BL/6 mouse, and six female 30-month-old Göttingen minipigs. Animal procedures were performed in accordance with the applicable Swedish, Israeli, or Swiss legislation on the protection of animals and were approved by the corresponding committees.

Murine experiments were approved by the local Swiss ethical committee (Tierversuchskommission des Kantons Bern, Amt für Veterinärwesen, Bern, Switzerland) under permit number BE 55/16. Minipig experiments were approved by the Malmö/Lund Ethics Committee on Animal Testing at the Lund District Court, Sweden under license 5.8.18-15672/2019. The following reporting adheres to the ARRIVE Guidelines 2.0 [39] for relevant items. Animal model, strain, age, animal number and sex are collected in a table available in the <u>Supplementary Materials</u>.

### Tomographic imaging

For this study, we imaged all samples with different Bruker SkyScan X-ray microcomputed tomography scanners (all from Bruker microCT N.V., Kontich, Belgium). An overview of the imaging and reconstruction parameters is given in the text below. Imaging parameters are mentioned briefly below and summarized in detail in a table given in the <u>Supplementary Materials</u>.

### Animal preparation and perfusion of contrast agent μAngiofil

The vasculature of all animals was instilled with the iodine-based contrast agent μAngiofil (Fumedica AG, Switzerland).

The contrast agent μAngiofil was prepared following the manufacturer’s instructions by first combining the two components ‘Contrast Solution’ and ‘PU’. These pre-mixed components were then mixed with a hardener immediately before instillation into the cannulated vascular system of the animals. Perfusion was conducted using a syringe pump to maintain consistent flow rates between 1 and 1.5 ml/min, ensuring reproducibility across experiments.

The perfusion of the animals was performed as previously described [21,22]. Heparinized animals were deeply anesthetized. Details on anesthesia for each animal model are provided below.

For mice, the descending aorta was cannulated, and blood was flushed with PBS before perfusion with μAngiofil. The exposed aorta was then cannulated in either antegrade (for the perfusion of the hind limbs) or retrograde direction (for the perfusion of the head and teeth) with a Venflon cannula (26 GA). The perfusion lasted until the organ of interest appeared completely filled with the blue contrast agent [20,22]. In bones, it is not possible to visually monitor this color change, thus perfusion of the neighboring soft tissues serves as an indirect marker of sufficient perfusion within the bone. To achieve correct perfusion of the vessels within the bone, we prolonged the perfusion time by instilling at least 2 ml of extra volume of contrast agent after all the superficial tissues of the extremity or head turned blue.

For minipigs, the external carotid artery was cannulated, and perfusion was performed similarly.

### Contrast-Enhanced microangioCT of Murine Tibia

Here, one 21-month-old VEGF transgenic male mouse was anesthetized with a mixture of fentanyl (0.05 mg/kg), midazolam (5 mg/kg), and medetomidine (0.5 mg/kg). μAngiofil perfusion was performed as described above. After μAngiofil polymerization, the sample was fixed in 4 % paraformaldehyde (PFA) solution at 4 °C and stored in PFA until tomographic imaging.

The sample was removed from the PFA solution, wrapped in closed-pore foam, and scanned in a custom-made sealed plastic sample holder under humid conditions using a SkyScan 1172. Imaging parameters included an acceleration voltage of 49 kV, a current of 200 μA, and an isotropic voxel size of 2.99 μm. We acquired 7202 projection images, recorded over a sample rotation of 360° (4000 x 2672 px, 3 averaged to one, each exposed for 985 ms).

Figure 1 shows the tibial bone microstructure and vascularization of a 21-month-old VEGF transgenic male mouse. This visualization approach enables the simultaneous display of bone and its vascularization. These tomographic datasets were also used in another study [14], where we performed simultaneous quantification of vasculature and bone volume.

**Figure 1:**
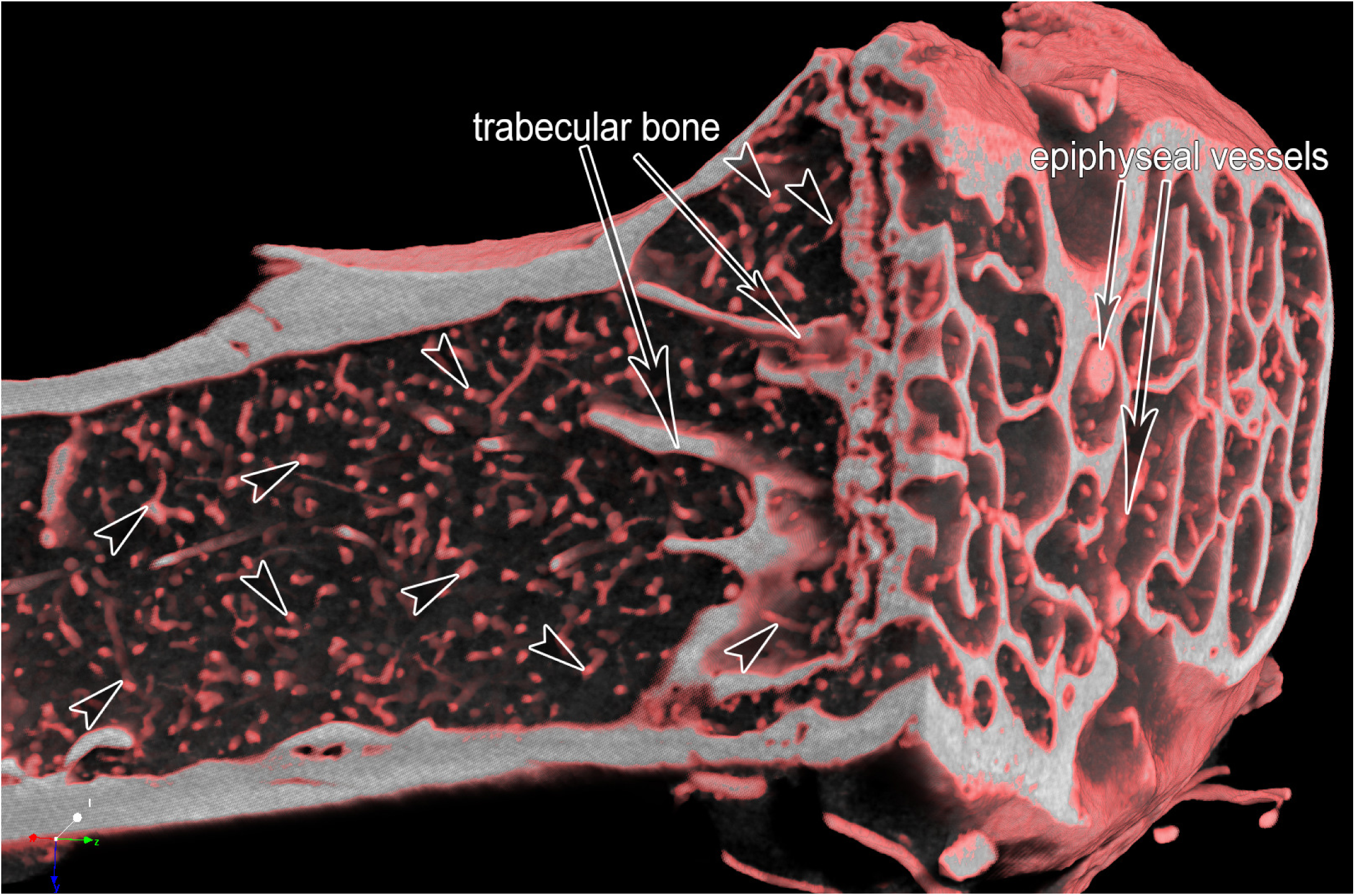
MicroangioCT of the proximal murine tibia of a 21-month-old VEGF transgenic male mouse. After perfusion with μAngiofil, the murine tibia was harvested, fixated in 4 % PFA and imaged by microCT. Arrowheads mark microvessels within the tibia. The diameter of the tibia shaft is around 1 mm. On the right side of the image, one can distinguish the bigger epiphyseal vessels. The bone tissue appears white at the plane of the virtual section through the microCT-dataset.

### Decalcification of the μAngiofil-perfused murine tibia

We established a bone decalcifying protocol using 10% ethylenediamine tetra-acetic acid (EDTA) solution [46], adapted from previous work [47].

Decalcification reduces the X-ray absorption of bone tissue without negatively influencing bone structure. However, this process renders the bone structure itself undetectable in tomographic imaging, preventing the simultaneous visualization of bone and vasculature (Fig. 2 B).

**Figure 2:**
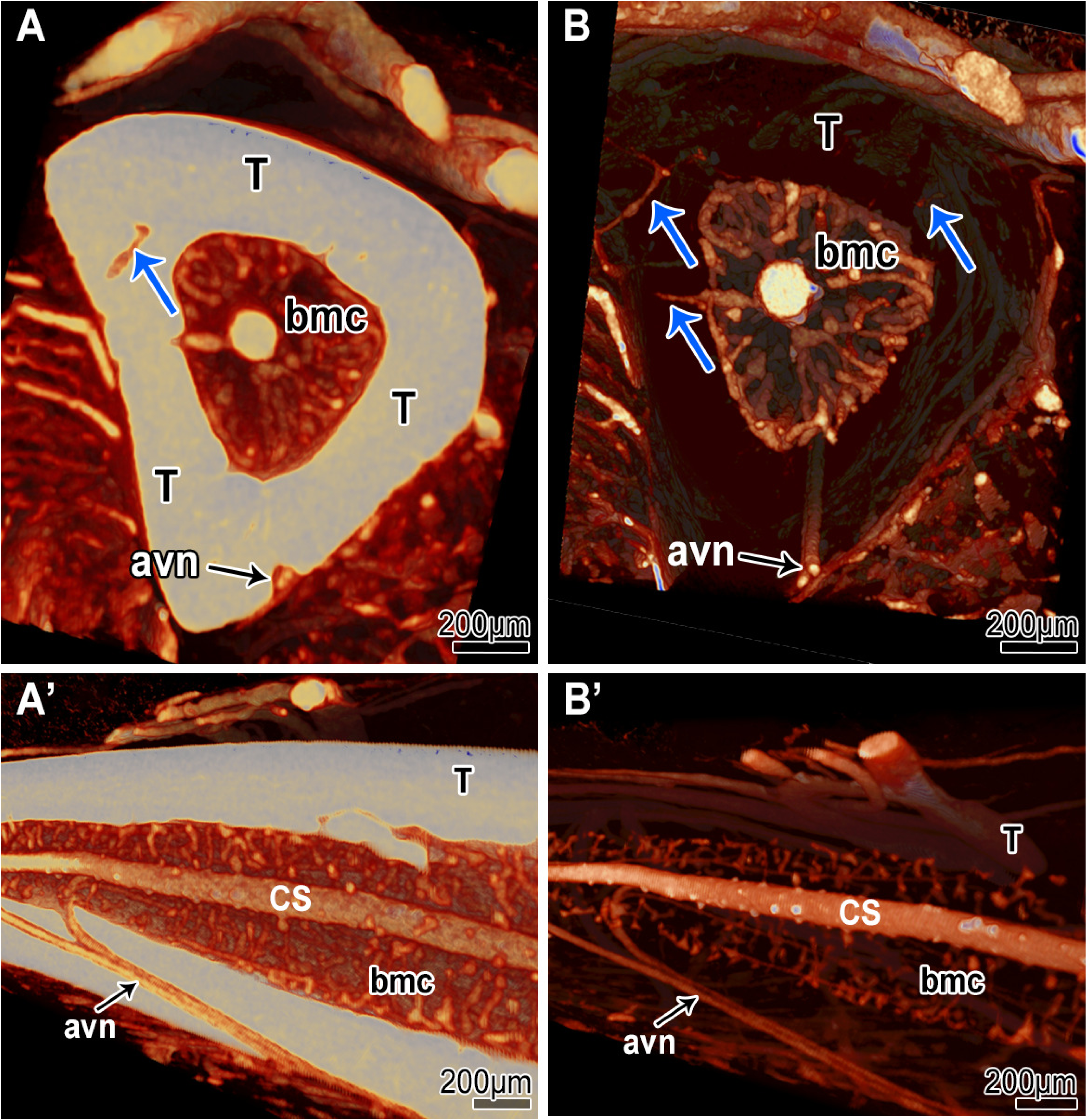
MicroangioCT-based visualization of the diaphysis of CB17SCID mice tibia before (A & A’) and after decalcification with 10% EDTA (B & B’). In A and A’ the tibia bone appears brighter and opaque due to higher X-ray absorption. In B and B’ the tibia bone appears transparent due to its lowered X-ray absorption after decalcification. Due to the decalcification, connecting vessels between the periosteal vessels and the vessels of the bone marrow cavity (bmc) are more easily detectable (blue arrows in A vs. B). The visualization of the vessels within the medullar cavity (*central sinus* (CS)) is also improved. At the external surface of the tibia, supplying arteries are visible (*arteria et vena nutricia* (avn)). The structure of the bone tissue is no longer clearly detectable after the decalcification (B-B’).

To evaluate the effects of decalcification, hind limbs of 10-week-old CB17SCID male mice were scanned before and after decalcification in 10% EDTA for 7 days.

Imaging parameters included an acceleration voltage of 59 kV, a current of 167 μA, and an isotropic voxel size of 3.19 μm. We acquired 1991 projection images, recorded over a sample rotation of 180°, (4000 x 2672 px, 2 averaged to one, each exposed for 1740 ms).

### Bone Metastatic Disease Model

In bone metastatic disease models, simultaneous imaging of bone and vasculature is crucial. Since changes in bone and vasculature are believed to be of crucial importance for the progression of bone metastatic disease, decalcification is generally avoided in these studies.

To verify the suitability of our microangioCT approach for imaging intratumoral vasculature in native, non-decalcified bone tissue we used the following murine bone metastatic disease model. 50000 PC3-M-Pro4Luc2 dTomato cells were injected into the tibia of 6-week-old CB17SCID male mice as previously described [8,56]. The X-ray assessment (25 kV, 6 sec, Faxitron Bioptics, Tucson, Arizona, US) was conducted on days 7, 14, 21, and 28 after implantation to monitor the progression of the lesions. Before perfusion with μAngiofil (as described above), the hind limb of interest was X-rayed using Faxitron Bioptics as a standard follow-up in this model (see insert in Fig. 3, Panel A). The harvested and fixated murine hind limb (4 % PFA at 4 °C) was then imaged using a SkyScan 1272 (Fig. 3). Imaging parameters included an acceleration voltage of 60 kV, a current of 166 μA, and an isotropic voxel size of 1.65 μm. We acquired 1878 projection images, recorded over a sample rotation of 180°, (4904 x 3280 px, 3 averaged to one, each exposed for 2800 ms). The visualization (CTvox (v.3.3.1), Bruker microCT N.V., Kontich, Belgium) displays intratumoral vasculature and extensive defects in mineralized bone tissue. Neighboring structures, such as the growth plate and the epiphysis or the menisci are also easily assessed (Fig. 3).

**Figure 3:**
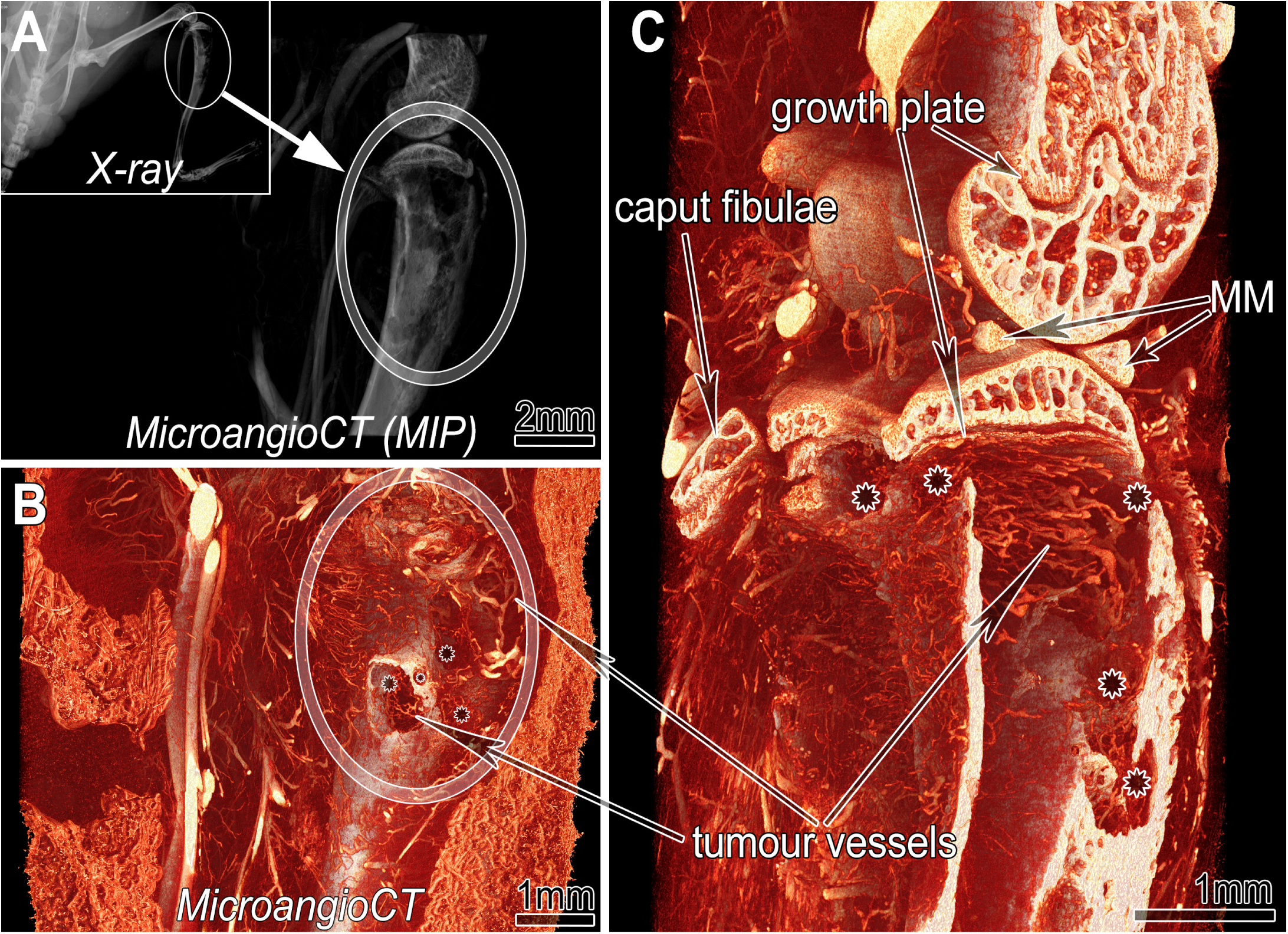
MicroangioCT of a xenograft tumor implanted into tibia of 6-week-old CB17SCID male mice. A: Maximum intensity projection (MIP) of the three-dimensional dataset of the investigated hind limb segment indicated in the inserted X-ray image of the mouse prior to harvesting. Inset: X-ray image before sample harvesting. B: Virtual section through the microangioCT dataset visualizing remarkable defects represented as holes (asterisks in B and C) in the tibial bone at the tumor site (encircled). C: A more deeply positioned virtual section displaying the inner surface of the diseased tibia. Besides irregularly patterned tumor vessels (in B & C) further bony structures like the growth plate or calcified parts of the medial meniscus (MM) are clearly distinguishable.

### Imaging of Murine Mandible and Teeth

C57BL/6 mice were anesthetized (with a mixture of fentanyl (0.05 mg/kg), midazolam (5 mg/kg), and medetomidine (0.5 mg/kg)), and their head was perfused with μAngiofil. The perfused head was harvested and fixed in 4 % PFA at 4 °C. Before tomographic imaging the mandible was excised, wrapped in a paper towel, and scanned in a sealed pipette tip.

The mouse teeth were imaged using a SkyScan 1172 (Fig. 4). Imaging parameters included an acceleration voltage of 80 kV, a current of 124 μA, and an isotropic voxel size of 1 μm. We acquired 3602 projection images, recorded over a sample rotation of 360°, (4000 x 2672 px, 4 averaged to one, each exposed for 6260 ms).

**Figure 4:**
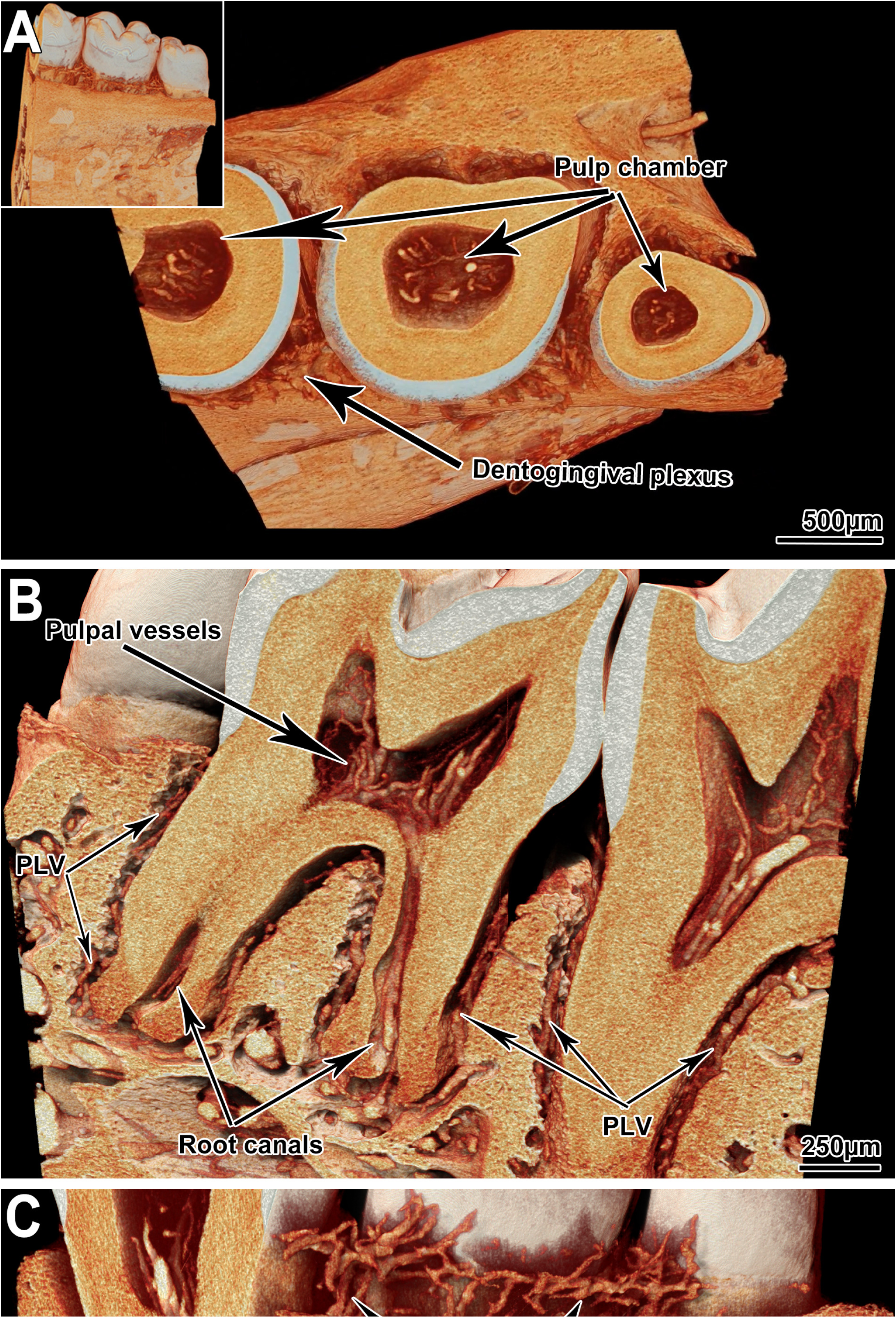
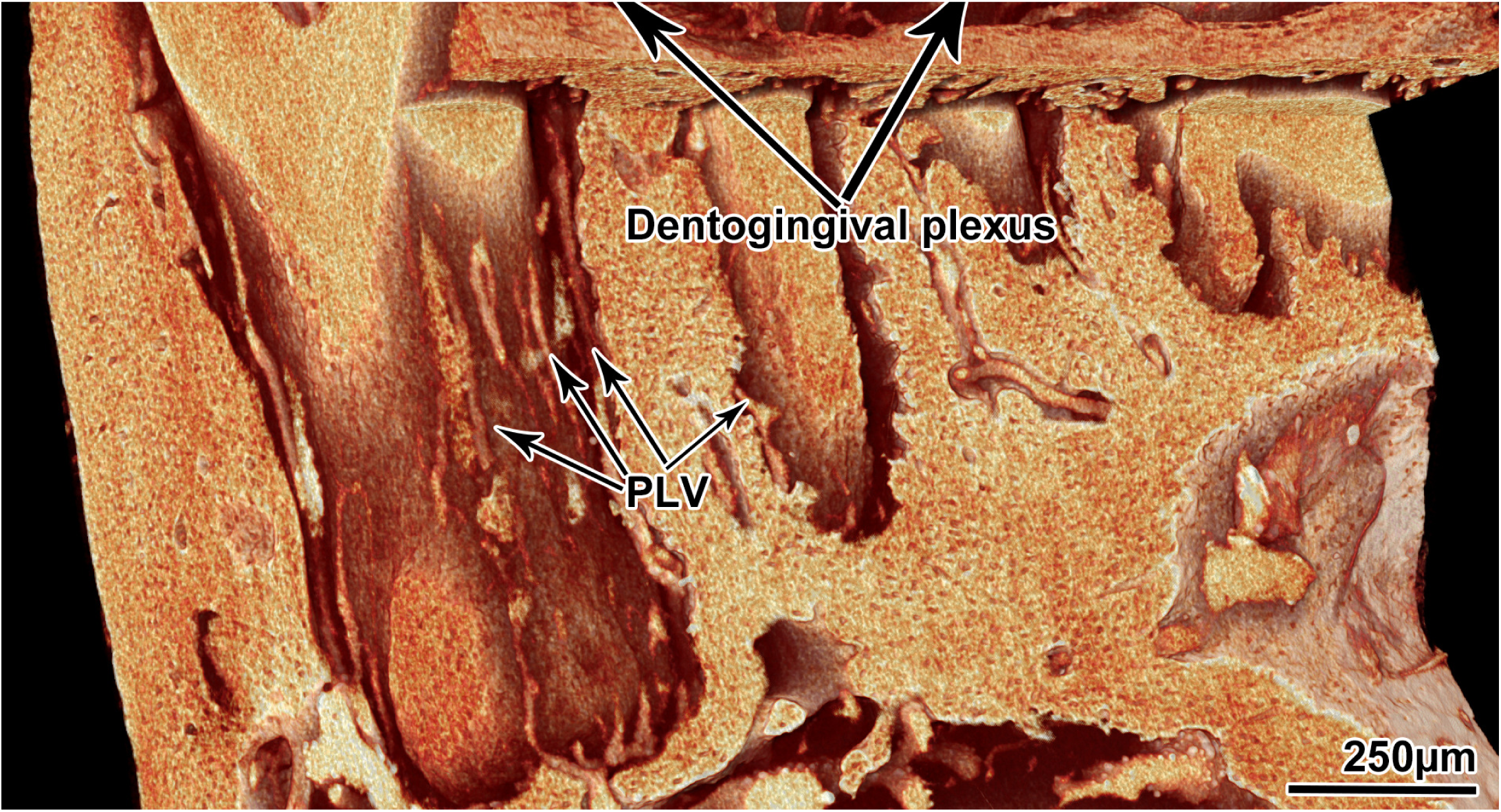
MicroangioCT of the vasculature of C57BL/6 mice teeth. A: View onto a virtual section parallel to the crowns of the murine teeth: pulp chambers are visible, and the pulpal vessels are presented. The inset shows a full view of the tomographic dataset. B: Sagittal section through the mandible. The microvessels within the pulp cavities and root canals are distinguishable. C: Detailed view of the dentogingival plexus and periodontal ligament vessels (PLV).

For murine mandibles and teeth, decalcification is unnecessary due to the distinct X-ray attenuation properties of μAngiofil and bone tissue. This allows clear visualization of the microvasculature of the vasculature within the mandible, periodontal ligament, and the teeth (even within their pulp chamber) without compromising bone structure (Fig. 4).

### Imaging of Minipig Mandible

Göttingen minipigs (Ellegaard Göttingen Minipig, Dalmose, Denmark) were anesthetized intramuscularly (25–35 mg/kg, Dexdomitor; Orion Pharma Animal Health and 50–70 mg/kg, Zoletil 100 Vet, Virbac) and intravenously heparinized with 300 IE/kg (Heparin LEO, LEO Pharma). After heparin infusion, the pigs were euthanized with an intravenous dose (100 mg/kg) of pentobarbital (Euthanimal vet, VM Pharma). The external carotid artery was accessed by blunt dissection through the tissue of the ventral neck and cannulated (BD Venflon, 17G). After washing out the blood with PBS, the corresponding head side was selectively perfused with μAngiofil through the arterial tree. Otherwise, the perfusion was performed with the same approach as in the previously described mice experiments. After the polymerization of μAngiofil, we excised the mandible and fixated and stored it in 4 % PFA solution at 4 °C.

Mandibles were then scanned with a SkyScan 1273 (Fig. 5). Imaging parameters included an acceleration voltage of 100 kV, a current of 80 μA, and an isotropic voxel size of 21 μm (panel A) or 9 μm (panels B and C). For the visualization shown in panel A, we recorded 3601 projection images over a sample rotation of 360° (3072 x 1944 px, 5 averaged to one, each exposed for 225 ms). For the visualizations shown in panels B and C, we recorded 2401 projection images over a sample rotation of 360° (5 averaged to one, each exposed for 225 ms). In addition, two horizontally overlapping projections were stitched to one projection with a size of 4832 x 1944 px to increase the imaged sample volume and cover the full extent of the sample.

**Figure 5:**
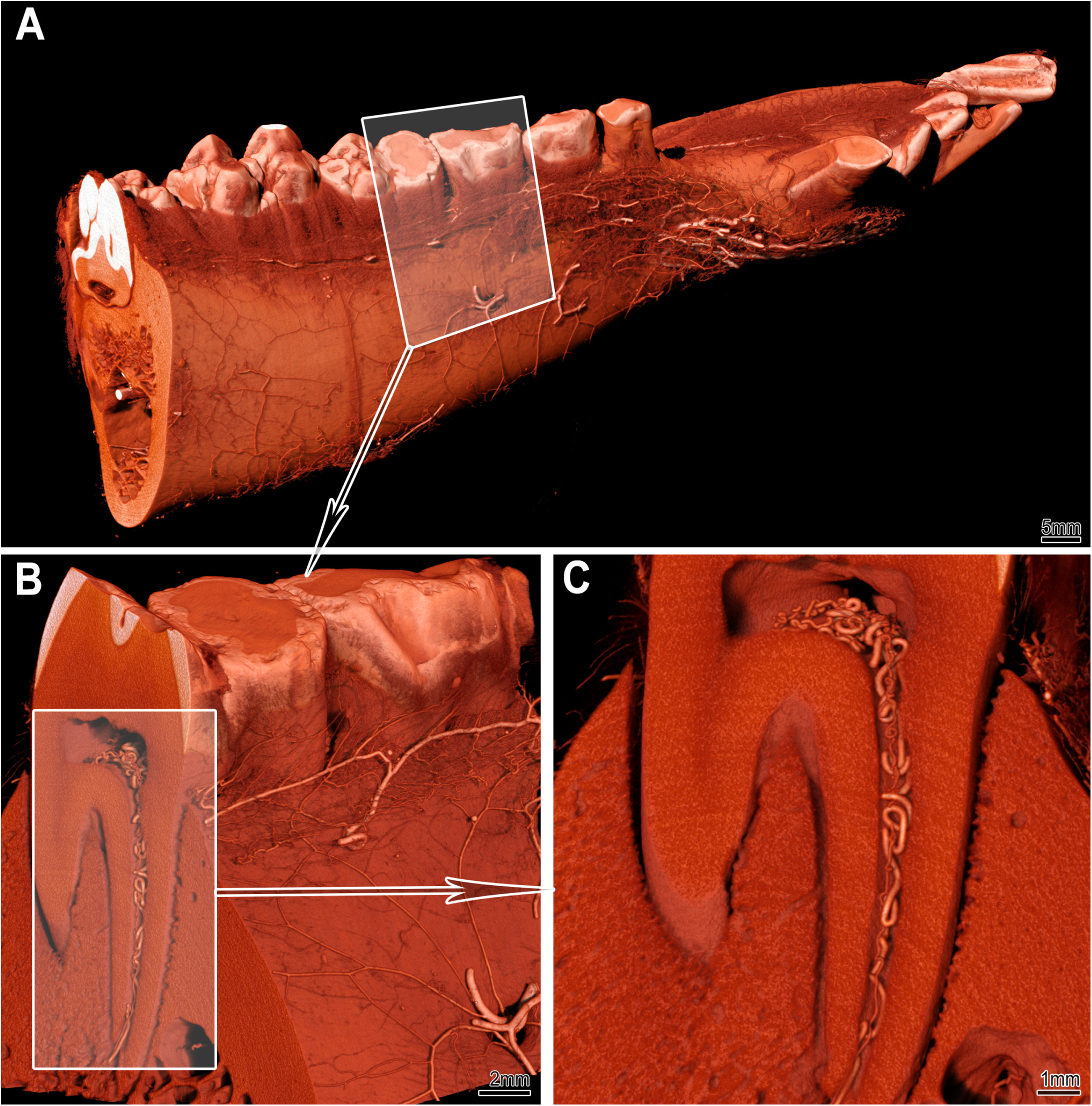
MicroangioCT of the minipig mandibula. Panel A displays the visualization of a right minipig hemimandible. The vasculature at the bone surface is visible. The framed area in A marks the subvolume represented in panel B at higher magnification. Panel C displays the transverse section marked in panel B, clearly showing the pulp chamber, root canal, and the corresponding vessels. Due to the voxel size of 8–9 μm, microvessels with a diameter of 40 μm or less cannot be visualized in such large samples.

Angiogenesis influences the osseointegration of implants. As such, studying angiogenesis and, correspondingly the vascular supply of the peri-implant tissue in detail is important for dental research and many implantology studies. So far, the only reliable approach to assess the vascular supply remains histology, limited to single two-dimensional sections.

While microCT imaging allows for non-destructive, full 3D imaging of dental research samples with implants, such imaging presents challenges. This is due to the presence of high-density metal parts with strong X-ray absorption within the samples. To distinguish the vasculature from both the metal implants and the mineralized bone tissue, we instilled the vasculature with a suitable contrast agent (μAngiofil). The X-ray absorption characteristics of μAngiofil make it possible to visualize and differentiate between soft tissue, bone tissue, contrast agent-filled vessels and metal (titanium) implants. We imaged an μAngiofil-instilled minipig hemimandibula with a SkyScan 2214 (Fig. 6), Imaging parameters included an acceleration voltage of 100 kV, a current of 100 μA, and an isotropic voxel size of 8 μm. We acquired 2001 projection images, recorded over a sample rotation of 360°, (2929 x 1944 px, 4 averaged to one, each exposed for 1080 ms).

**Figure 6:**
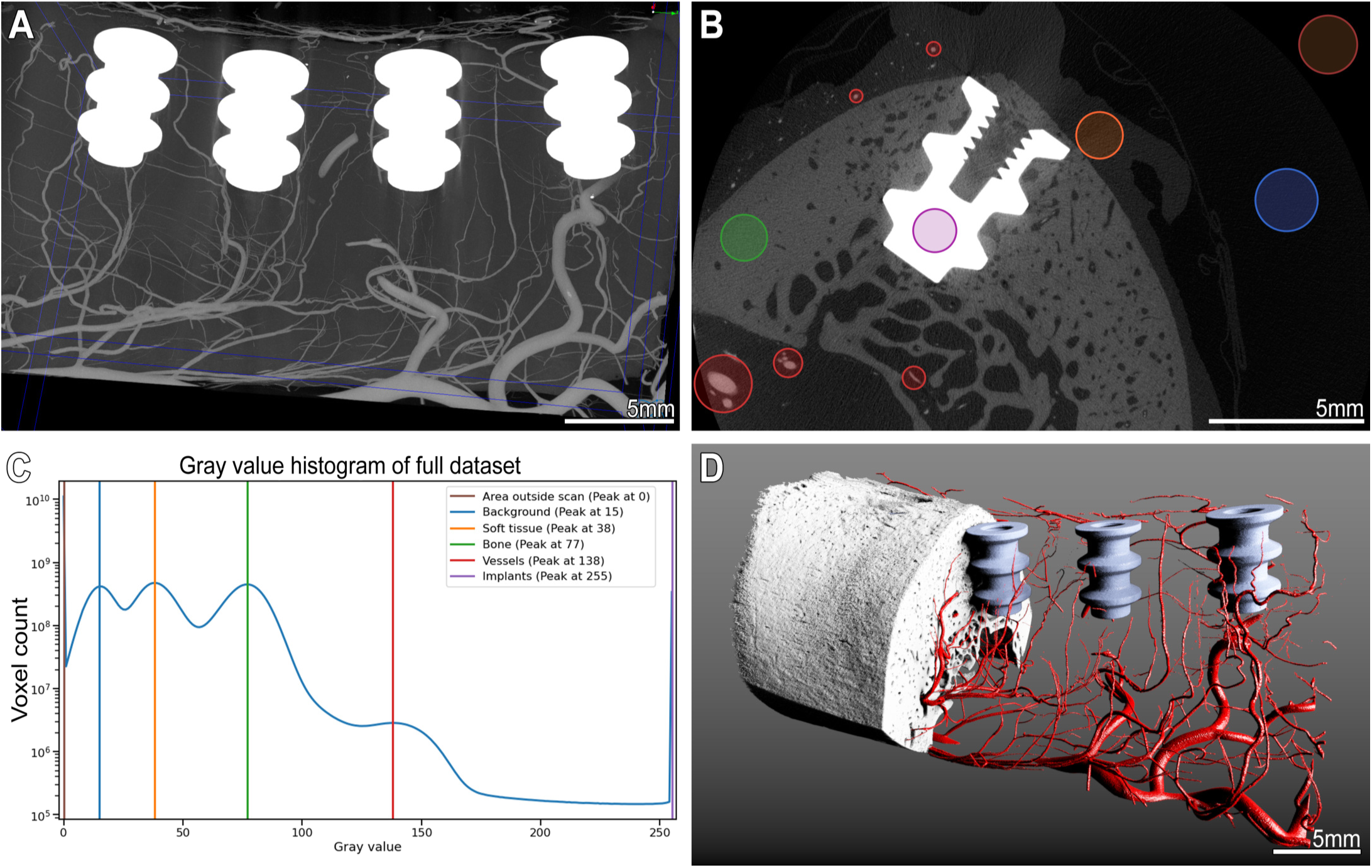
MicroangioCT of the peri-implant vasculature of a minipig mandibula. Panel A: Maximum intensity projection image of the minipig mandible dataset with four metal implants and μAngiofil-perfused vessels. Panel B: virtual transversal section through the dataset showing an implant within the mandible. The colored circles mark structures with different gray value ranges. Corresponding peaks with color legend are marked on the histogram in panel C. Such differences in gray levels allow a straightforward segmentation of the structures of interest as displayed in the 3D visualization in panel D.

In the resulting tomographic datasets these regions of interest can easily be distinguished based on their gray value ranges, as shown in Fig. 6, Panel C. Tomographic imaging of such samples and straightforward segmentation of features of interest without cumbersome post-processing (Fig. 6, Panel D) is enabled without requiring a decalcification step.

### Dual-Energy microangioCT

Unfortunately, not all samples exhibit a pronounced difference in gray values between bone tissue and μAngiofil when scanned using standard scanning parameters. This issue is particularly common in murine samples, even though the bone mineral density of mouse bones is generally higher than that of other experimental species [9,25]. In tomographic datasets of murine samples, the gray values of bone tissue and μAngiofil are similar making it impossible to distinguish them based on histogram levels alone (e.g., Fig. 4). While decalcification can improve contrast (see Fig. 2), we propose a more efficient alternative: dual-energy scanning Previous studies have demonstrated that microscopic dual-energy CT imaging (microDECT) using commercial lab-based microCT systems can achieve spectral separation of two or more materials with micrometer resolution [15,16]. Notably, the contrasting properties of hydroxyapatite (the primary mineral in bone and teeth) and iodine (the high-Z component in μAngiofil) make them ideal for spectral X-ray imaging [11,15,28].

We evaluated multiple energies (accelerating voltages from 40 to 110 kV) to choose the optimal combination for imaging of the murine mandible perfused with μAngiofil using a SkyScan 2214 (Fig. 7). The optimal dual-energy settings for the X-ray source were 50 kV/120 μA and 90 kV/100 μA, respectively (acceleration voltage/source current). We acquired 3601 projection images, recorded over a sample rotation of 360° (4032 x 2688 px, 7 averaged to one, each exposed for 4550 ms). This resulted in datasets with an isotropic voxel size of 1.4 μm.

**Figure 7:**
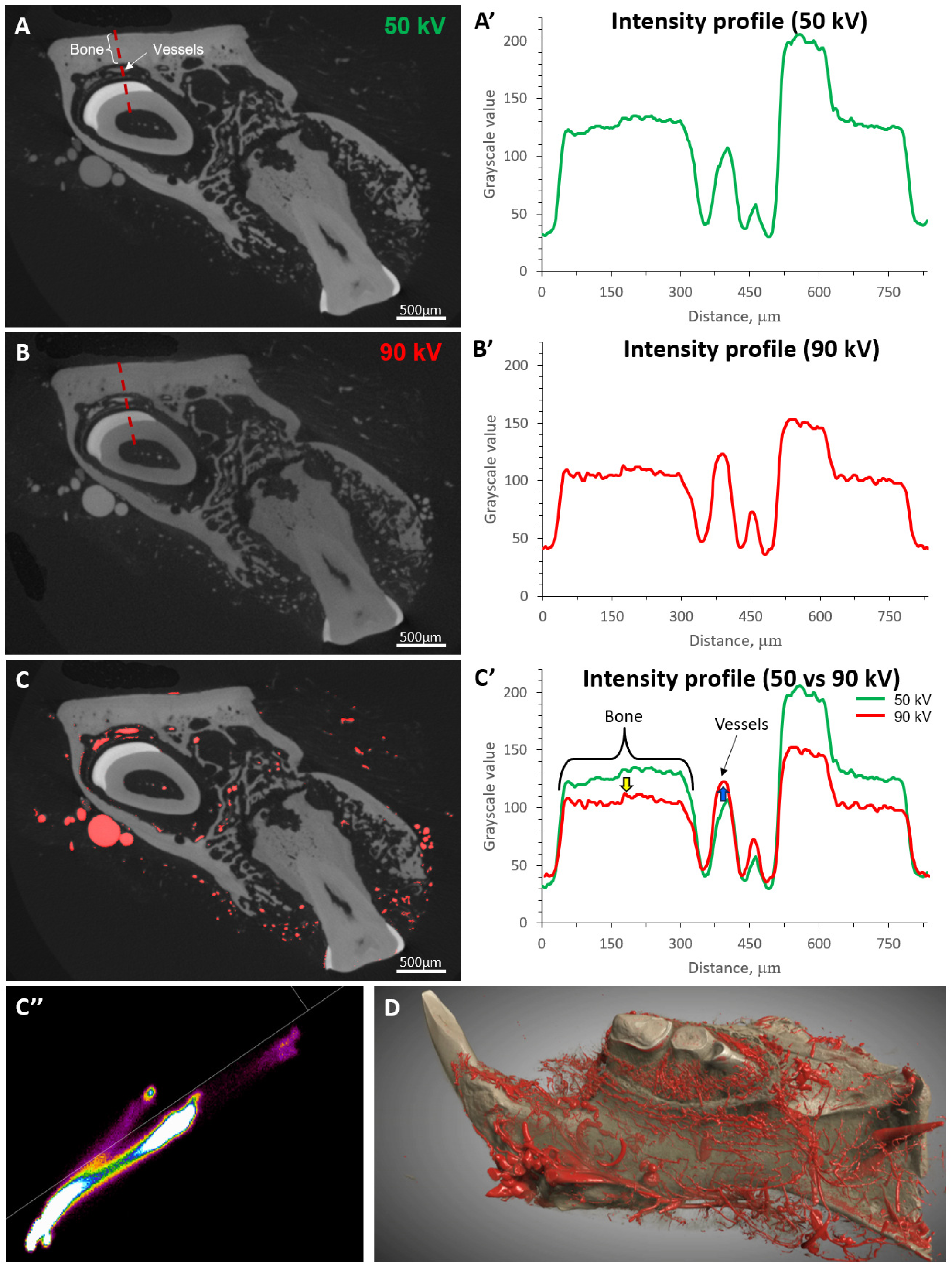
Dual-energy segmentation of murine mandible vasculature. Panel A and B represent virtual sections through the microCT datasets obtained at different accelerating voltages: 50 kV (A) and 90 kV (B). The dashed red line in panels A and B represents the location of the corresponding intensity profile displayed in panels A’ and B’. Panel C highlights the vasculature which is segmented with using the dual-energy approach in red. The changes in the intensity profiles of different tissues are shown in graph C’: with the higher voltage (90 kV) the intensity of bone tissue reduces (yellow arrow) and the intensity of vessels (μAngiofil) increases. Such changes in gray levels allow segmentation of the vessels using the combined histogram shown in panel C’’ (DEhist, Bruker microCT N.V., Kontich, Belgium). Panel D represents a three-dimensional visualization of the segmented tissues (Dragonfly 3D World (2024.1), Comet Technologies Canada Inc., Montréal, Canada).

In comparison to 50 kV, at 90 kV the gray value of the bone tissue was decreased and the gray value of μAngiofil increased (Fig. 7 C’). By using different X-ray energies, this method enhances contrast between bone and μAngiofil without compromising bone structure, enabling straightforward segmentation based on the combined histogram (Fig. 7 C’’).

## Discussion

Adequate vascularization is a prerequisite for successful bone formation and regeneration as well as osseointegration of biomaterials [6,32,48,49,52]. Without high-resolution imaging, the interplay between vasculature and bone tissue cannot be adequately studied, making it impossible to control these processes.

Non-destructive microCT imaging is widely regarded as the only viable method for 3D imaging of an intact bone-implant interface [54]. To accurately differentiate vasculature from other soft tissue structures, the use of a contrast agent or the creation of a vascular replica is essential.

The imaging method we present, which involves instillation with μAngiofil, addresses several limitations of previously described techniques (see <u>Introduction</u>). This approach enables the visualization of both bone tissue and the microvasculature within it, with or without prior decalcification (see Fig. 2).

The established decalcifying protocols enable correlative imaging of the same sample with preserved intravascular contrast agent with subsequent classical histological evaluation [46]. This decalcification step enables tomographic imaging of the murine hind limb vasculature with fewer artifacts around the bone at lower X-ray source acceleration voltages due to the reduced radiopacity of the sample.

Lower acceleration voltages often allow for shorter scanning times, leading to higher sample throughput. Furthermore, decalcifying the sample enables a threshold-based segmentation of the vasculature, and facilitates histological evaluation of the sample, as previously described [46].

However, the decalcification step renders bone tissue transparent, preventing simultaneous 3D imaging of the bony and vascular structures.

In many models, due to the distinct differences in the X-ray attenuation of μAngiofil and the bone tissue, we can forgo the time-consuming decalcification step. This permits us direct assessment of vascularization and bone growth in a single scan (Fig. 6). While effective, this approach does not always yield optimal results (e.g., Fig. 4). Although decalcification is a viable option for quite some studies, a dual-energy scanning technique offers a more efficient solution (see Fig. 7). By conducting two sequential scans without removing the sample, this method simplifies dataset registration, avoids the time-consuming decalcification step, and provides simultaneous imaging of bone and vasculature while keeping the sample intact. Therefore, the microangioCT approach we present here can be used for the simultaneous visualization of hard and soft tissues and their vascularization in small and large animal models.

One of the noteworthy applications of this approach is the simultaneous visualization of the bone microarchitecture and microvasculature of bone metastases in a murine xenograft tumor model (Fig. 3). Especially for studying bone metastatic models, it is crucial to find a method that allows for the correct assessment of the microvasculature without decalcifying the sample. Decalcification would hinder the thorough assessment of pathological processes in bone tissue. These pathological processes can cause skeletal-related events, which are associated with shortened survival and deterioration of quality of life [7], and should therefore be avoided. Our method allows studying the response to specific treatments, including bone-targeted and antiangiogenic therapies. These can be assessed in 3D within the same sample and followed up with a histological examination if desired [53]. The presented method could offer additional insights into the interplay between angiogenesis, bone growth, bone lysis, and bone turnover, which have not been fully elucidated yet. The potential findings can be crucial for selecting potential drug candidates, making proper treatment decisions, and efficacy prediction.

In dental research, preclinical models can be divided into small and large animals. Small animal models, particularly mice and rats, are highly popular, due to their practical size and cost-effectiveness. The microCT scan of correspondingly small samples, perfused with μAngiofil can be performed with a voxel size of around 1 μm, providing excellent detail resolution (see Fig. 4). Murine mandibles and teeth are challenging for assessing the vasculature. This is due to the location of most of the vessels within the bone canals or in the proximity of the hard tissue. This leads to a lack of larger bone-free volumes where the vasculature is easily distinguishable. Gray value-based segmentation of bone and contrast-agent-instilled vasculature is compounded by the fact that murine bone has a higher mineral bone density when compared to other species [9,25]. Nonetheless, with the dual-energy approach described above, we achieved appropriate imaging of such samples and were able to visualize and distinguish vasculature within and from mineralized bone tissue.

Further work, also from our group, and ongoing research is focusing on improving the segmentation of vasculature in microtomographic datasets. Convolutional neural networks (CNNs) have emerged as a powerful tool for vascular segmentation [50]. Specifically, the U-Net architecture of CNNs seems very promising for vascular segmentation tasks and has been applied for the segmentation of murine vascular networks [36]. While neural network-based segmentation methods have shown promising results, the scarcity of annotated public datasets poses a challenge for training robust models and developing models that can generalize across different tissues, modalities, scales, and pathologies remains an active area of investigation.

In the small animal model studies without metal or similar biomedical implants, the application of whole mouse clearing and imaging with vDISCO approach could be the method of choice [4]. Another promising improvement to the clearing protocols and immunolabeling of the samples with the bone tissue is the introduction of the collagenase digestion step, which improves the antibody penetration throughout the stained bones and, therefore, the immunostaining [3]. Although newer tissue clearing and immunostaining immersion-based visualization techniques provide high-resolution, three-dimensional images of intact samples, these methods may produce artifacts such as tissue deformation and illumination inhomogeneity [45].

The presence of implanted biomedical devices further complicates the imaging process. For murine model studies, a recently published tissue-clearing-based imaging approach [54] could be a viable option for the visualization of the peri-implant tissues and vasculature due to the small size of hemimandible samples and the availability of transgenic mouse lines. However, there are limitations to such a tissue-clearing imaging approach. Namely: i) differential shrinkage among soft and hard tissues, leading to anisotropic distortion in samples where both tissue types (plus eventual metal implant) are present; ii) limited imaging depth (approximately 800 μm in a mouse model [54]) due to the challenges of achieving complete transparency of bone tissue. Significant anisotropic distortion in the sample may alter the implant-tissue interface, a common site of interest. The limitation of the maximal achievable imaging depth makes such an approach impractical for usage in large animal models.

Our presented microangioCT approach does not have such limitations and can be easily applied for visualization of hemimandible and its vascularization in a large animal model like Göttingen minipig (Fig. 5). Iodine-based μAngiofil exhibits significantly different attenuation properties than mineralized bone in most species and allows for the distinction, segmentation, and visualization of soft tissue, bone tissue, vessels filled with the contrast agent, and metal implants according to their gray values in the histogram (see Fig. 6).

To the best of our knowledge, our study is the first to demonstrate non-destructive 3D imaging of the microvasculature of bone in the proximity of metal objects/implants in a large animal model.

While this study primarily focused on qualitative visualization of vascular structures, we acknowledge the importance of quantitative analyses. Future work will incorporate metrics such as vessel diameter, connectivity, and perfusion efficiency to provide a more comprehensive evaluation of µAngiofil’s performance.

## Limitations of our approach

Achieving the excellent results demonstrated in this study requires reproducible and controlled instillation of the contrast agent. To ensure this, the procedure should be performed by skilled personnel using a syringe pump. While other contrast agents may be easier to use for instillation, they do not deliver the same high-quality results as previously discussed.

The method we presented for investigating the microvasculature within bone tissue is primarily limited by the imaging technique. Specifically, this limitation arises from the correlation between achievable resolution and the physical size of the imaged sample. For small samples, such as those just a few millimeters in diameter, voxel sizes on the order of 1 μm enable visualization of the microvasculature down to the capillary bed [20–22,46]. However, for larger samples, such as the minipig hemimandible, the achievable voxel size is approximately 10 times larger (≥8 μm). In such cases, the finest microvessels with diameters under 40 μm cannot be accurately visualized (see Fig. 5 & <u>6</u>).

Beyond the inherent limitations of tomographic imaging, the approach presented here—instilling the vasculature in bone tissue with a polymerizing contrast agent—represents a significant advancement in biomedical X-ray imaging. This innovative method holds great promise for addressing key questions in tissue engineering, implantology, and a wide range of related research fields.

## Author Contributions

Contributor Roles Taxonomy, as defined by the National Information Standards Organization.

| Author | Contributions |
| --- | --- |
| David Haberthür | Data curation; Formal analysis; Investigation; Project administration; Software; Validation; Visualization; Writing – original draft; Writing – review & editing |
| Oleksiy-Zakhar Khoma | Data curation; Formal analysis; Investigation; Project administration; Software; Validation; Visualization; Writing – original draft; Writing – review & editing |
| Tim Hoessly | Data curation; Investigation; Writing – review & editing |
| Eugenio Zoni | Investigation; Methodology; Resources; Writing – review & editing |
| Marianna Kruithof-de Julio | Methodology; Resources; Writing – review & editing |
| Stewart D. Ryan | Resources; Writing – review & editing |
| Myriam Grunewald | Investigation; Methodology; Resources; Writing – review & editing |
| Benjamin Bellón | Investigation; Methodology; Writing – review & editing |
| Rebecca Sandgren | Investigation; Writing – review & editing |
| Stephan Handschuh | Methodology; Writing – review & editing |
| Benjamin E. Pippenger | Funding acquisition; Investigation; Methodology; Resources; Writing – review & editing |
| Dieter Bosshardt | Methodology; Writing – review & editing |
| Valentin Djonov | Conceptualization; Funding acquisition; Resources; Writing – review & editing |
| Ruslan Hlushchuk | Conceptualization; Funding acquisition; Investigation; Methodology; Project administration; Resources; Supervision; Validation; Visualization; Writing – original draft; Writing – review & editing |

## Conlicts of interest

Conflicts of interest
| Author | Conflict |
| --- | --- |
| Benjamin Bellón | Employed by a company producing dental implants |
| Benjamin E. Pippenger | Employed by a company producing dental implants |

## Acknowledgments

We are grateful to the Microscopy Imaging Center of the University of Bern for infrastructural support.

We thank the project [18] for helping us write this manuscript collaboratively.

## Supplementary Materials

### Log files of all the tomographic scans performed for this study

The table shown in <u>SampleAndScanData.csv</u> gives a tabular overview on the samples studied and the relevant parameters of the tomographic imaging. This file was generated with a <u>data processing</u> <u>notebook</u>and contains information read from *all* the log files of *all* the scans we performed. A copy of each log file is also available <u>online</u>. These log files include *all* the data necessary to exactly replicate the image acquisition.

## Data availability

Tomographic datasets used and analyzed during the current study is available from the corresponding author upon reasonable request.

